# CB_1_ receptor-mediated inhibitory LTD triggers presynaptic remodeling via protein synthesis and ubiquitination

**DOI:** 10.1101/2020.01.09.900464

**Authors:** Hannah R. Monday, Mathieu Bourdenx, Bryen A. Jordan, Pablo E. Castillo

**Affiliations:** Dominick P. Purpura Department of Neuroscience, Albert Einstein College of Medicine, 1300 Morris Park Avenue, Bronx, NY, 10461, USA; Department of Developmental and Molecular Biology, Albert Einstein College of Medicine, 1300 Morris Park Avenue, Bronx, NY, 10461, USA; Institute for Aging Studies, Albert Einstein College of Medicine, 1300 Morris Park Avenue, Bronx, NY, 10461, USA; Department of Psychiatry and Behavioral Sciences, Albert Einstein College of Medicine, 1300 Morris Park Avenue, Bronx, NY, 10461, USA

**Keywords:** structural plasticity, presynaptic, protein synthesis, ubiquitination

## Abstract

Long-lasting forms of postsynaptic plasticity commonly involve protein synthesis-dependent structural changes of dendritic spines. However, the relationship between protein synthesis and presynaptic structural plasticity remains unclear. Here, we investigated structural changes in cannabinoid-receptor 1 (CB_1_)-mediated long-term depression of inhibitory transmission (iLTD), a form of presynaptic plasticity that requires protein synthesis and involves a long-lasting reduction in GABA release. We found that CB_1_-iLTD in acute rat hippocampal slices was associated with protein synthesis-dependent presynaptic structural changes. Using proteomics, we determined that CB_1_ activation in hippocampal neurons resulted in increased ribosomal proteins and initiation factors, but decreased levels of proteins involved in regulation of the actin cytoskeleton, such as Arp2/3, and presynaptic release. Moreover, while CB_1_-iLTD increased ubiquitin/proteasome activity, ubiquitination but not proteasomal degradation was critical for structural and functional presynaptic CB_1_-iLTD. Thus, CB_1_-iLTD relies on both protein synthesis and ubiquitination to elicit structural changes that underlie long-term reduction of GABA release.

## INTRODUCTION

Synaptic plasticity, the ability of synapses to change their strength in response to activity or experience, underlies information storage in the brain. While presynaptic forms of plasticity, i.e. long-term synaptic strengthening (long-term potentiation or LTP) and weakening (long-term depression or LTD) due to long-lasting increase and decrease in neurotransmitter release, respectively, are widely expressed in the brain, their mechanism remains poorly understood (Castillo, 2012; Monday & Castillo, 2017; Monday et al., 2018; Yang & Calakos, 2013). A good example of a ubiquitous form of long-lasting reduction of transmitter release in the brain is type-1 cannabinoid receptor (CB_1_)-mediated LTD (Araque et al., 2017; Castillo et al., 2012; Heifets & Castillo, 2009). Here, endogenous cannabinoids (eCBs) are released upon activity and travel in a retrograde manner to bind presynaptic CB_1_, a G_i/o_-coupled receptor, resulting in CB_1_-LTD at both excitatory and inhibitory synapses. Induction of long-term eCB-mediated plasticity requires extended (minutes) CB_1_ activation (Chevaleyre & Castillo, 2003; Chevaleyre et al., 2007; Ronesi et al., 2004). Although the presynaptic changes downstream CB_1_ that suppress transmitter release in a long-term manner remain unclear, there is evidence that presynaptic protein synthesis is required (Yin et al., 2006; Younts et al., 2016), but what proteins are synthesized and the precise role of these proteins in CB_1_-LTD remain unexplored.

Proteostatic mechanisms, the cellular processes that balance protein synthesis and degradation, are vital for neuronal function and synaptic plasticity (Biever et al., 2019; Birdsall & Waites, 2019; Cohen & Ziv, 2019; Liang & Sigrist, 2018; Y. C. Wang et al., 2017). In postsynaptic forms of plasticity, such as NMDA receptor-dependent LTP, local protein synthesis has been tightly correlated with both consolidation of LTP and structural changes (Bosch et al., 2014; Ostroff et al., 2002; Tanaka et al., 2008; Tominaga-Yoshino et al., 2008; Yang et al., 2008). In particular, synthesis of β-actin, AMPA receptors, and CaMKII proteins may be critical for the increase in dendritic spine volume and synapse strength associated with LTP (Bramham, 2008; Nakahata & Yasuda, 2018; Rangaraju et al., 2017). Concurrent regulation of protein degradation through the proteasome and lysosome is also required for activity-dependent pre- and postsynaptic changes in synapse strength (Biever et al., 2019; Cohen & Ziv, 2017; Hegde, 2017; Monday et al., 2018). We and others have recently provided evidence for rapid (<30 min) presynaptic protein synthesis under basal conditions and during plasticity (Hafner et al., 2019; Younts et al., 2016), but whether these newly synthesized proteins participate in CB_1_-LTD by regulating presynaptic structural changes is unknown.

Presynaptic structure and function are controlled by actin polymerization and depolymerization (Cingolani & Goda, 2008; Nelson et al., 2013). Branched actin filaments in the presynaptic compartment provide a scaffold for synaptic vesicles and the active zone (Michel et al., 2015; Rust & Maritzen, 2015). Moreover, structural changes of the presynaptic terminal are associated with altered synapse strength (Gundelfinger & Fejtova, 2012; Matz et al., 2010; Meyer et al., 2014; Monday & Castillo, 2017). There is evidence that CB_1_ activation alters the ultrastructural vesicle distribution in CB_1_-expressing (CB_1_^+^) boutons on short time scales (Garcia-Morales et al., 2015; Ramirez-Franco et al., 2014) and leads to retraction of growth cones in developing axons (Roland et al., 2014). However, it is not known whether CB_1_-LTD is associated with morphological changes in presynaptic structure in the mature mammalian brain.

Here, to gain insights into the expression mechanisms of CB_1_-LTD, we examined potential structural changes in CB_1_-mediated LTD of inhibitory transmission (CB_1-_iLTD) in the hippocampus (Chevaleyre & Castillo, 2003). Using high-resolution microscopy in acute rat hippocampal slices we found that this form of plasticity was associated with a reduction of presynaptic bouton volume that required protein synthesis. To test how protein synthesis could alter presynaptic structure during CB_1_-iLTD, we used an unbiased proteomics approach to identify CB_1_ activation-mediated changes in the proteome of hippocampal neuron cultures. CB_1_ activation elicited an increase in proteins involved in protein synthesis, processing and degradation, whereas presynaptic and actin cytoskeletal proteins, including Arp2/3 and WAVE1 underwent rapid degradation. CB_1_-iLTD involved actin remodeling, Rac1 and the actin branching protein Arp2/3. Lastly, ubiquitination of proteins but not proteasomal degradation was necessary for both structural and functional CB_1_-iLTD.

## RESULTS

### Induction of CB_1_-iLTD is associated with a reduction in in CB_1_+ bouton size

Diverse forms of long-lasting synaptic plasticity require translation-dependent structural remodeling (Bailey et al., 2015; Bramham, 2008; Nakahata & Yasuda, 2018; Rangaraju et al., 2017). To test whether CB_1_-iLTD is associated with structural plasticity, we examined changes in the morphology of CB_1_^+^ boutons following induction of CB_1_-iLTD. To accurately measure individual bouton volume in acute hippocampal slices, we utilized high-resolution and high yield Airyscan confocal microscopy combined with 3D reconstruction (**Figure 1A**). CB_1_-iLTD was induced by bath application of the CB_1_ agonist WIN 55,212-2 (25 min, 5 µM) (Heifets et al., 2008; Younts et al., 2016) (**Figure 1B**). This LTD not only mimics synaptically-induced iLTD (Chevaleyre & Castillo, 2003; Chevaleyre et al., 2007; Heifets et al., 2008), but also allows us to shortcut the synthesis and release of eCBs, thereby excluding potential effects of pharmacological inhibitors (see below) on these processes.

**Figure 1:**
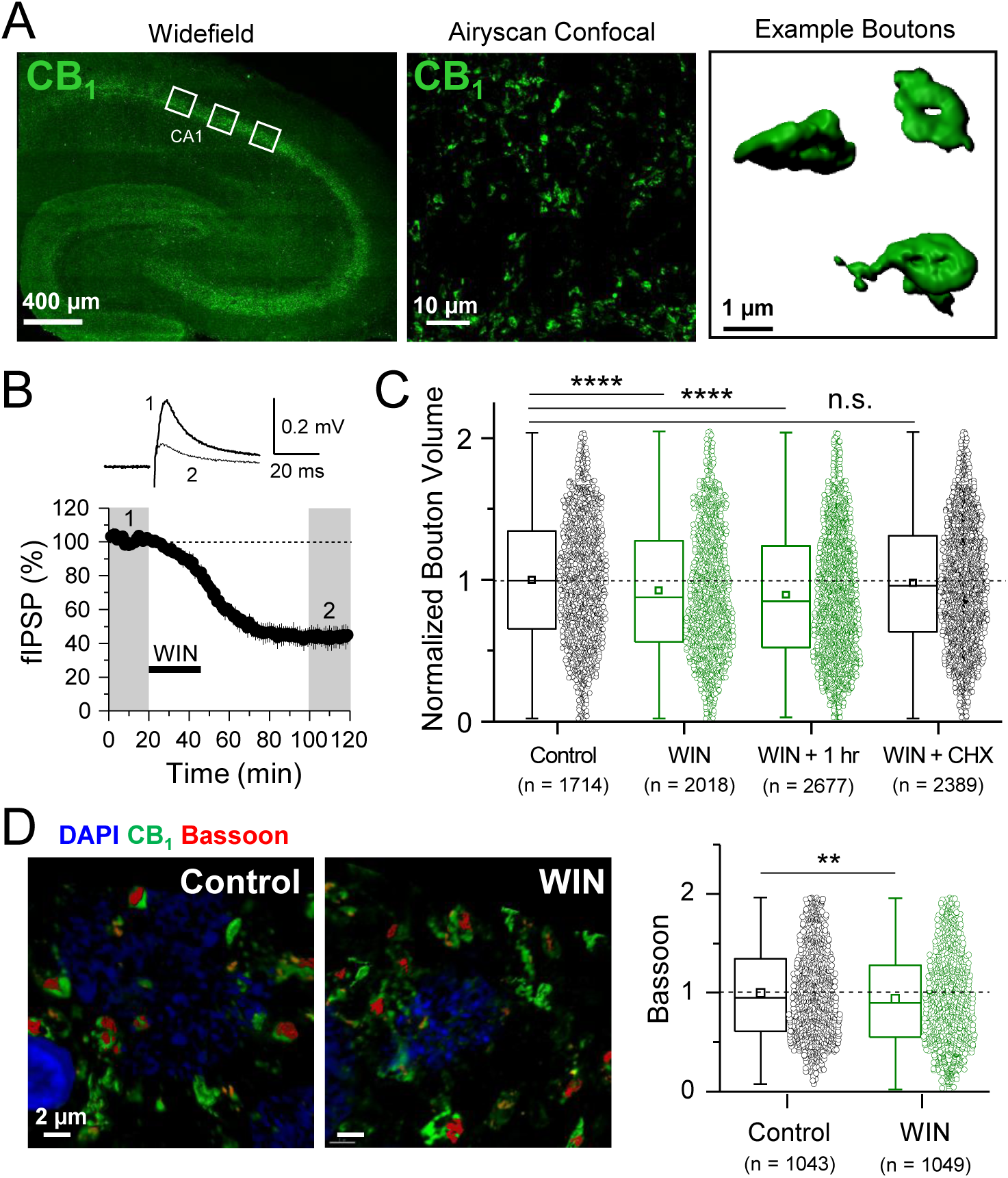
Induction of CB_1_-iLTD is associated with a reduction in CB_1_^+^ bouton size. *A. Left*, representative widefield confocal of hippocampal CB_1_ immunolabeling. White boxes indicate where high magnification images were acquired (as seen in center panel). *Center*, High-resolution Airyscan confocal maximum projection used for 3D reconstruction of individual boutons in CA1 stratum pyramidale. *Right*, representative single boutons reconstructed in 3D using Imaris. *B. Top*, representative extracellular field inhibitory postsynaptic potentials (fIPSPs) recorded in the CA1 pyramidal cell body layer in acute hippocampal slices before and after WIN treatment (5 µM, 25 min). *Bottom*, summary time-course plot showing WIN-induced depression; n = 3 slices, 3 animals. C. Quantification of bouton volume normalized to control. Activation of CB_1_ receptors with 5 µM WIN for 25 min led to decreased bouton volume that remained 1 hour after WIN treatment and was blocked by treatment with cycloheximide (CHX, 80 µM, applied throughout the experiment). Control: 1.0 ± 0.01 vs. WIN: 0.92 ± 0.01 vs. WIN + 1 hr: 0.89 ± 0.01 vs. CHX + WIN: 0.98 ± 0.01; F[4,10770] = 21.17; **** indicates p < 0.0001; n.s., non significant; one-way ANOVA. n = number of boutons (3 images/slice, 2 slices/rat, 6 rats/condition). For all structural plasticity figures, data are presented as box plots (left) and data points (right) where box represents the 25^th^ and 75^th^ percentile of data range, mean is represented with a square, and median with a line inside the box. Minimum and maximum of data are given by the capped line. *D. Left*, representative 3D reconstruction images of Bassoon labeling inside CB_1_^+^ boutons. *Right*, quantification of Bassoon volume after 25 min WIN treatment revealed a reduction in active zone volume as measured using Bassoon immunolabeling. Control: 1.0 ± 0.01 vs. WIN: 0.94 ± 0.02, U = 585539.5, Z = 2.79, ** indicates p<0.01, Mann-Whitney, n = number of active zones in CB_1_^+^ boutons (2 images/slice, 2 slices/rat, 5 rats/condition).

Using CB_1_ immunolabeling, which accurately approximates bouton volume (Dudok et al., 2015), we found that CB_1_-iLTD led to a significant decrease of CB_1_ bouton volume following WIN treatment (**Figure 1C**). This structural change was long-lasting as it persisted for 60 minutes after WIN treatment (**Figure 1C**), and it was blocked by concurrent bath application with cycloheximide (80 µM), demonstrating a requirement for protein synthesis (**Figure 1C**). The effect was specific because the volume of parvalbumin (PV^+^) boutons in the CA1 pyramidal layer, which do not express CB_1_ receptors (Glickfeld & Scanziani, 2006) (**Supp. Figure 1A**), was not altered by WIN application (**Supp. Figure 1B,C**), as assessed by PV immunolabeling (Younts et al., 2016). As a complementary approach independent of CB_1_ labeling, we used Bassoon immunolabeling to assess the size of the presynaptic active zone. Similar to the total bouton volume, Bassoon size within CB_1_^+^ boutons was also significantly decreased following WIN application (**Figure 1D**). These results strongly suggest that CB_1_-iLTD is associated with a protein synthesis-dependent shrinkage of CB_1_^+^ boutons, which may contribute to the long-lasting reduction in neurotransmitter release observed in this form of plasticity. Along with our previous study (Younts et al., 2016), our findings indicate that protein synthesis is required for both structural and functional presynaptic changes involved in CB_1_-iLTD.

### CB_1_ activation alters the abundance of proteins linked to protein synthesis, synaptic structure/function and energy metabolism

To glean insights into the mechanism(s) underlying structural and functional CB_1_-iLTD, we sought to identify proteins synthesized upon CB_1_ activation. We previously showed CB_1_-dependent increases in protein synthesis were evident after brief CB_1_ activation in cultured hippocampal neurons (Younts et al., 2016). To identify and quantitate changes in the neuronal proteome, we used Stable Isotope Labeling of Amino Acids in Cell Culture (SILAC) coupled with tandem mass spectrometry (MS/MS) (Jordan et al., 2006; G. Zhang et al., 2011; G. Zhang et al., 2012). Two populations of cultured rat hippocampal neurons (‘medium’ and ‘heavy’) were labeled using distinct combinations of stable-isotope variants of Arginine and Lysine. The two groups were treated with WIN (25 min, 5 µM) as before or Vehicle (**Figure 1**) then rapidly lysed and harvested **(Supp. Figure 2A, B)**. Samples were combined and simultaneously analyzed by tandem MS/MS to identify and quantify changes induced by CB_1_ receptor activation. To strengthen the robustness of findings, we performed a replicate “reverse” experiment where ‘heavy’ neurons were treated with WIN and observed a high degree of correlation between replicates (**Supp. Figure 2C**).

We found significant changes across the protein landscape. Setting a threshold cutoff at 0.5 log_2_-fold change, we found that 33 proteins were upregulated and 27 proteins were downregulated by CB_1_ activation. Examples of these proteins grouped by their suggested function are shown in **Figure 2A (see Supp. Table 1 for all proteins)**. Consistent with previous studies of axonal mRNAs, components of the protein synthesis machinery were upregulated (Hafner et al., 2019; Shigeoka et al., 2016), as well as the protein degradation machinery. A number of presynaptic proteins were downregulated following CB_1_ activation. Notably, two key regulators of the actin cytoskeleton, Actin-related protein 2/3 (Arp2/3) and Wiscott-Aldrich Associated Protein family (WAVE1) were significantly downregulated, suggesting these proteins could be implicated in the reduction of neurotransmitter release and presynaptic volume associated with CB_1_-iLTD. Using Gene Set Enrichment Analysis (GSEA) (Subramanian et al., 2005), we identified enriched functional gene ontology (GO) terms (**Figure 2B and Supp. Table 2**). To reduce redundancy, we clustered closely related GO terms using network analysis (Merico et al., 2010), where edge length corresponds to the number of overlapping proteins in the GO term, node size indicates the number of proteins belonging to the term, and color represents the enrichment score (**Figure 2C and Supp. Table 2**). In accordance with translational upregulation following CB_1_ activation (Younts et al., 2016), the top cluster represented upregulated GO terms related to “Protein synthesis and processing” (**Figure 2C**). The second cluster was composed of GO terms relating to “Neuronal projections”, suggestive of the structural change associated with CB_1_-iLTD. The third cluster was GO terms associated with “Energy metabolism” which may be representative of the previously reported CB_1_-mediated decrease in cellular respiration (Hebert-Chatelain et al., 2016; Mendizabal-Zubiaga et al., 2016) (**Figure 2C**). Examples of GO terms identified in each cluster are provided in **Supp. Figure 3**. Ingenuity Pathway Analysis (IPA) also identified pathways related to the processes outlined above, including eIF2 signaling, Mitochondrial Dysfunction, and Actin Cytoskeleton Signaling **(Supp. Figure 2D).** Similarly, analyses using SynGO (Koopmans et al., 2019), an expert-curated tool to identify GO terms associated with synaptic function, linked our results to regulation of synaptic protein synthesis **(Supp. Figure 2E).** Together, these results suggest that both protein synthesis and coincident degradation of structural and presynaptic proteins occur downstream of CB_1_ activation, and could therefore be implicated in CB_1_-iLTD.

**Figure 2:**
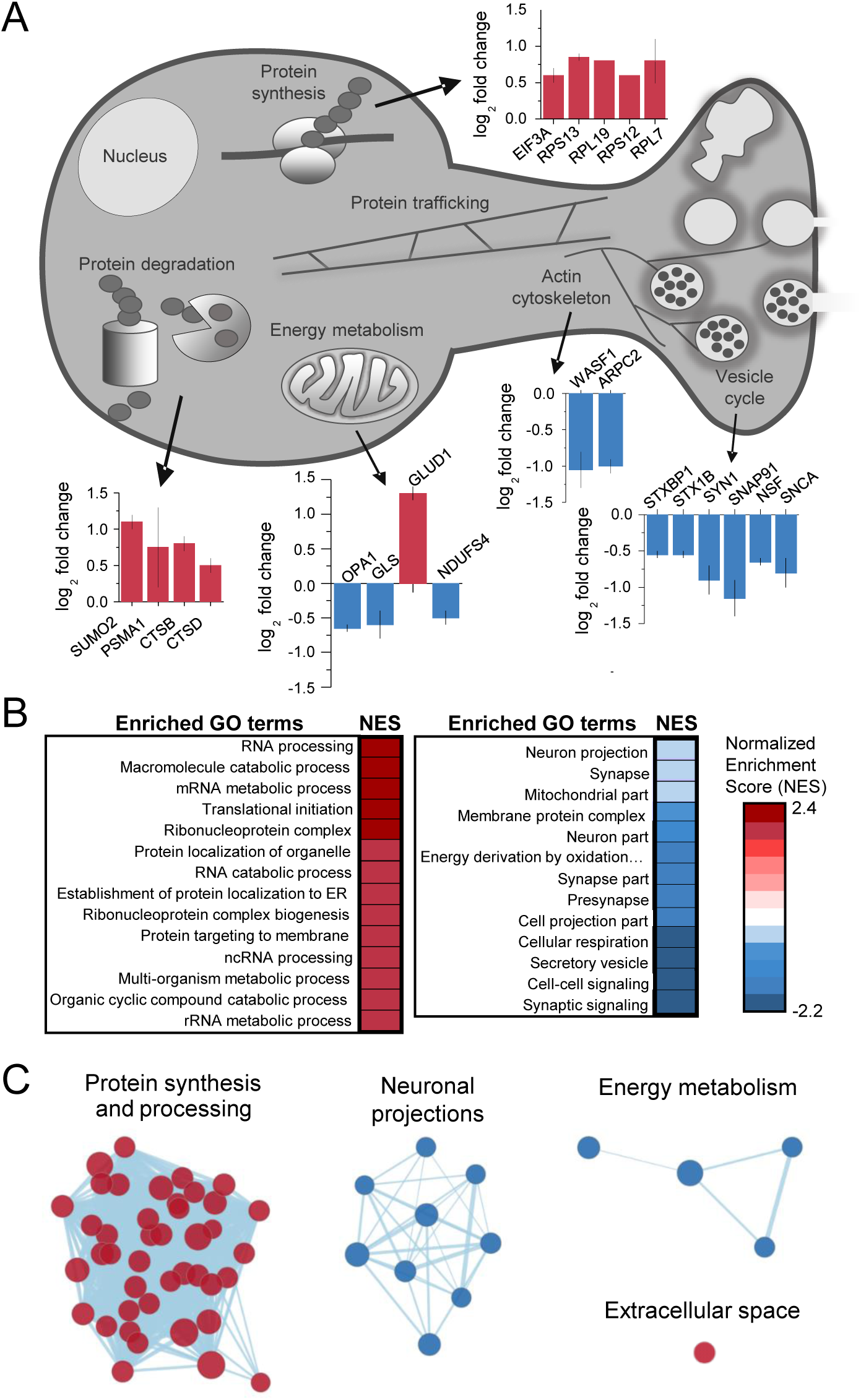
CB_1_ activation alters the abundance of proteins linked to protein synthesis, synaptic structure/function and energy metabolism. A. Examples of proteins that were identified in enriched GO terms and were significantly altered by CB_1_ activation (p < 0.05). Protein IDs are grouped by proposed biological function and average log_2_ fold change is plotted. B. List of enriched GO terms and normalized enrichment scores (NES) as identified by GSEA. Positive NES reflects overall upregulation of proteins associated with the GO term whereas negative values indicate the opposite. C. Cluster analysis of enriched/depleted GO terms from GSEA revealed 4 distinct biological processes that were consistently up or downregulated by CB_1_ activation. Each node represents a single GO term. Node size represents magnitude of enrichment and edge length gives degree of overlap between 2 GO terms. Color represents up (red) or downregulation (blue) of proteins associated with that GO term. See Supp. Figure 3 for examples.

### CB_1_-iLTD involves actin remodeling via Rac1 and Arp2/3

CB_1_ directly interacts with actin branching modulators WAVE1 and Arp2/3 (Njoo et al., 2015), and these proteins are downregulated in hippocampal neurons following CB_1_ activation (**Figure 2A**), therefore regulation of the abundance of these proteins may represent a mechanism underlying structural and functional presynaptic changes involved in CB_1_-iLTD. For example, CB_1_ activation could reduce the presynaptic terminal volume by favoring actin depolymerization. To test this possibility, we first examined whether actin cytoskeletal dynamics were required for CB_1_-iLTD induced structural plasticity (**Figure 1**). Using the same high-resolution microscopy and 3D reconstruction as **Figure 1**, we activated CB_1_ in the presence of jasplakinolide (JSK, 250 nM), a reagent that promotes actin-stabilization (Holzinger, 2009). We found that JSK application blocked the WIN-induced decrease in presynaptic bouton volume (**Figure 3A**). These results indicate that actin dynamics likely underlie the structural changes following CB_1_ activation. To test the functional requirement for actin remodeling in CB_1_-iLTD, we monitored inhibitory synaptic transmission in the CA1 area by recording extracellular field inhibitory postsynaptic potentials (fIPSP) (**see Figure 1**), which allows non-invasive, stable long-term assessment of inhibitory synaptic transmission, and induced CB_1_-iLTD by bath applying the CB_1_ agonist WIN 55, 212-2 (25 min, 5 µM) (Heifets et al., 2008; Younts et al., 2016). Similar to the effects on structural plasticity, bath application of JSK impaired CB_1_-iLTD (**Figure 3B**), whereas JSK application alone had no effect on basal synaptic transmission (**Supp. Figure 4A**). These results strongly suggest that actin remodeling is critical for structural and functional CB_1_-iLTD.

**Figure 3:**
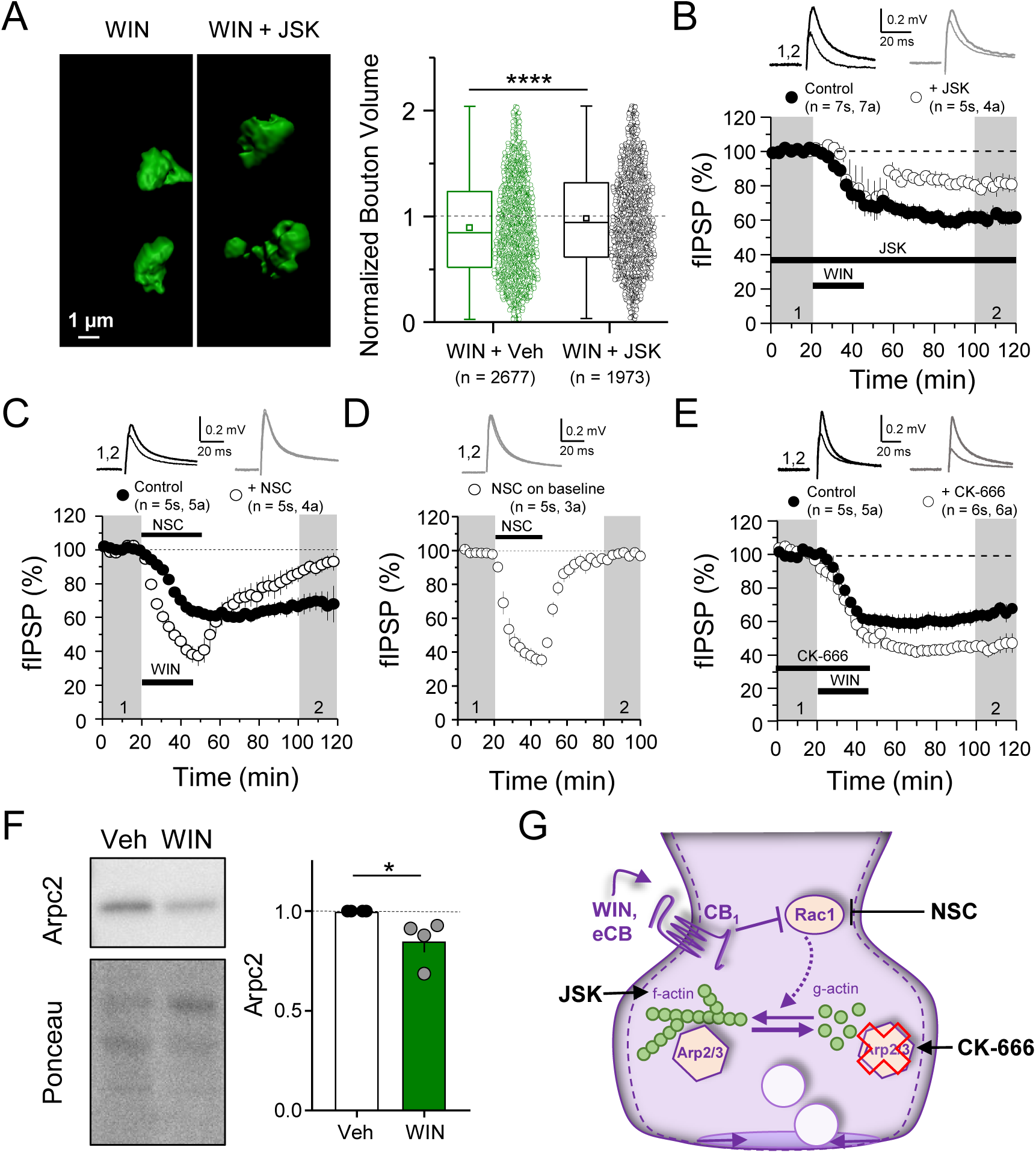
CB_1_-iLTD involves actin remodeling via Rac1 and Arp2/3. *A. Left*, representative single boutons reconstructed in 3D. *Right*, quantification of bouton volume normalized to control. Activation of CB_1_ receptors with WIN for 25 min led to decreased bouton volume that was blocked by treatment with jasplakinolide (JSK, 250 nM). Summary data expressed as normalized change from Vehicle ± S.E.M.. WIN + Veh: 0.89 ± 0.01 vs. JSK + WIN: 0.98 ± 0.01, U = 2.92E6, Z = 6.356, **** indicates p < 0.0001, Mann-Whitney, n = number of CB_1_^+^ boutons (3 images/slice, 2 slices/rat, 6 rats/condition). Data are presented as box plots (left) and data points (right) where box represents the 25th and 75^th^ percentile of data range, mean is represented with a square, and median with a line inside the box. Minimum and maximum of data are given by the capped line. B. CB_1_-iLTD is impaired by bath application of actin-stabilizing drug, jasplakinolide (JSK, 250 nM). Extracellular field inhibitory postsynaptic potential (fIPSP) were recorded in the CA1 pyramidal cell body layer in acute hippocampal slices. Control: 61.4 ± 4 % vs. JSK: 80.5 ± 4 %; p < 0.05, unpaired t-test. For all electrophysiology figures, representative traces correspond to the gray shaded areas and in the summary time-course plots (averaged summary data expressed as normalized change from baseline ± S.E.M.), and n = number of slices (s), number of animals (a) unless otherwise specified. Shaded boxes in all electrophysiology figures correspond to when plasticity was analyzed with respect to baseline and when representative traces were collected and averaged. C. CB_1_-iLTD was blocked by acute bath application of the Rac1 inhibitor NSC (30 µM). Control: 68.8 ± 6 % vs. NSC23766: 89.7 ± 4 %; p < 0.05, unpaired t-test. D. NSC (30 µM) bath application reversibly depressed basal transmission. NSC: 98 ± 2 %, one sample t-test, p > 0.05. E. CB_1_-iLTD is enhanced by acute bath application of the Arp2/3 inhibitor CK-666 (100 µM). Control: 63.8 ± 4 % vs. CK-666: 45.2 ± 4 %, p > 0.05, unpaired t-test. *F. Left*, representative Western blots of staining for Arpc2 and Ponceau loading control in vehicle or WIN treated hippocampal cultures. *Right*, Arpc2 was downregulated in hippocampal neuron cultures after CB_1_ activation with WIN (5 µM, 25 min). Arpc2 (Fold of Veh): 0.851 ± 0.06, U = 16, Z = 2.31, * indicates p < 0.05, Mann-Whitney. Dots represent individual values for 4 independent experiments. Data in the bar plot represent mean ± S.E.M. G. Model of CB_1_-iLTD pathway and points of regulation. CB_1_ activation leads to inhibition of Rac1 which regulates actin dynamics via Arp2/3. Arp2/3 loss leads to actin depolymerization and increased CB_1_-iLTD. Actin remodeling is required for CB_1_-iLTD. NSC inhibits Rac1-GEF interaction. CK-666 stabilizes the inactive conformation of Arp2/3, preventing it from binding actin filaments. JSK stabilizes actin filaments and promotes polymerization.

The Rac1 GTPase is one of the principal regulators of actin polymerization via WAVE1 and Arp2/3 activity (Derivery & Gautreau, 2010; Stradal & Scita, 2006). To test the role of these proteins, we inhibited Rac1 activity using NSC 23766 (NSC), an inhibitor of Rac1-GEF interaction (Gao et al., 2004). CB_1_-iLTD was impaired by application of NSC (30 µM, 25 min) during induction (**Figure 3C**). NSC alone transiently suppressed inhibitory transmission (**Figure 3D**), unlike excitatory transmission (Hou et al., 2014), and this effect was associated with a decrease in PPR (**Supp. Figure 4B**), suggesting Rac1 activity regulates GABA release. To directly test the role of Arp2/3 in CB_1_-iLTD we utilized CK-666 (100 µM), a compound that inhibits Arp2/3-mediated actin assembly by stabilizing the inactive conformation of Arp (**see Figure 3G**) (Basu et al., 2016; Hetrick et al., 2013). CK-666 bath application enhanced CB_1_-iLTD (**Figure 3E**) suggesting that Arp2/3 participates in CB_1_-iLTD. Unlike NSC, CK-666 had no effect on basal inhibitory transmission (**Supp. Figure 4C**), presumably because the inhibitor stabilizes the inactive (unbound) Arp2/3, but does not affect the Arp2/3 bound to actin filaments. Arpc2 protein (Arp2/3 complex subunit 2), an essential component of the Arp2/3 complex, was degraded upon CB_1_ activation in hippocampal neuron cultures (**Figure 3F**), suggesting that during normal CB_1_-iLTD, CB_1_ activation-mediated Rac1 inhibition leads to removal of Arp2/3 from actin branches, and the subsequent degradation of Arp2/3 (**Figure 3G**). The enhancement of CB_1_-iLTD by CK-666 application probably occurs because the unbound Arp2/3 that is not degraded following CB_1_ activation becomes inhibited and cannot maintain actin branches, thereby resulting in further depolymerization. Together, our findings suggest that Rac1 signaling and loss of Arp2/3 likely underlie the actin remodeling, which is likely required for functional and structural CB_1_-iLTD (**Figure 3G**).

### CB_1_-iLTD requires ubiquitination, but not degradation by the proteasome

The simplest interpretation of our findings is that CB_1_-induced degradation of Arp2/3 and WAVE1 led to decreased actin polymerization and reduced presynaptic bouton size (**see Figure 2**). To test the role of protein degradation in this process, we first confirmed that presynaptic proteins identified in the SILAC screen, Munc18-1, Synapsin-1, and α-Synuclein, were significantly reduced by WIN (25 min, 5 µM) in hippocampal cultures (**Figure 4A**), suggesting rapid protein degradation upon CB_1_ activation. To assess whether presynaptic proteins are degraded locally in acute hippocampal slices, we prevented anterograde and retrograde axonal transport by incubating slices in nocodazole (1 hr, 20 µM), an agent that depolymerizes axonal microtubules (Barnes et al., 2010; Younts et al., 2016). We found that CB_1_ activation with WIN reduced Synapsin-1 puncta intensity in CB_1_^+^ boutons despite blockade of axonal transport (**Figure 4B**), as measured by immunostaining and quantitative Airyscan microscopy, consistent with local degradation. These results suggest that CB_1_ activation elicits rapid degradation of presynaptic proteins in culture and in acute slices.

**Figure 4:**
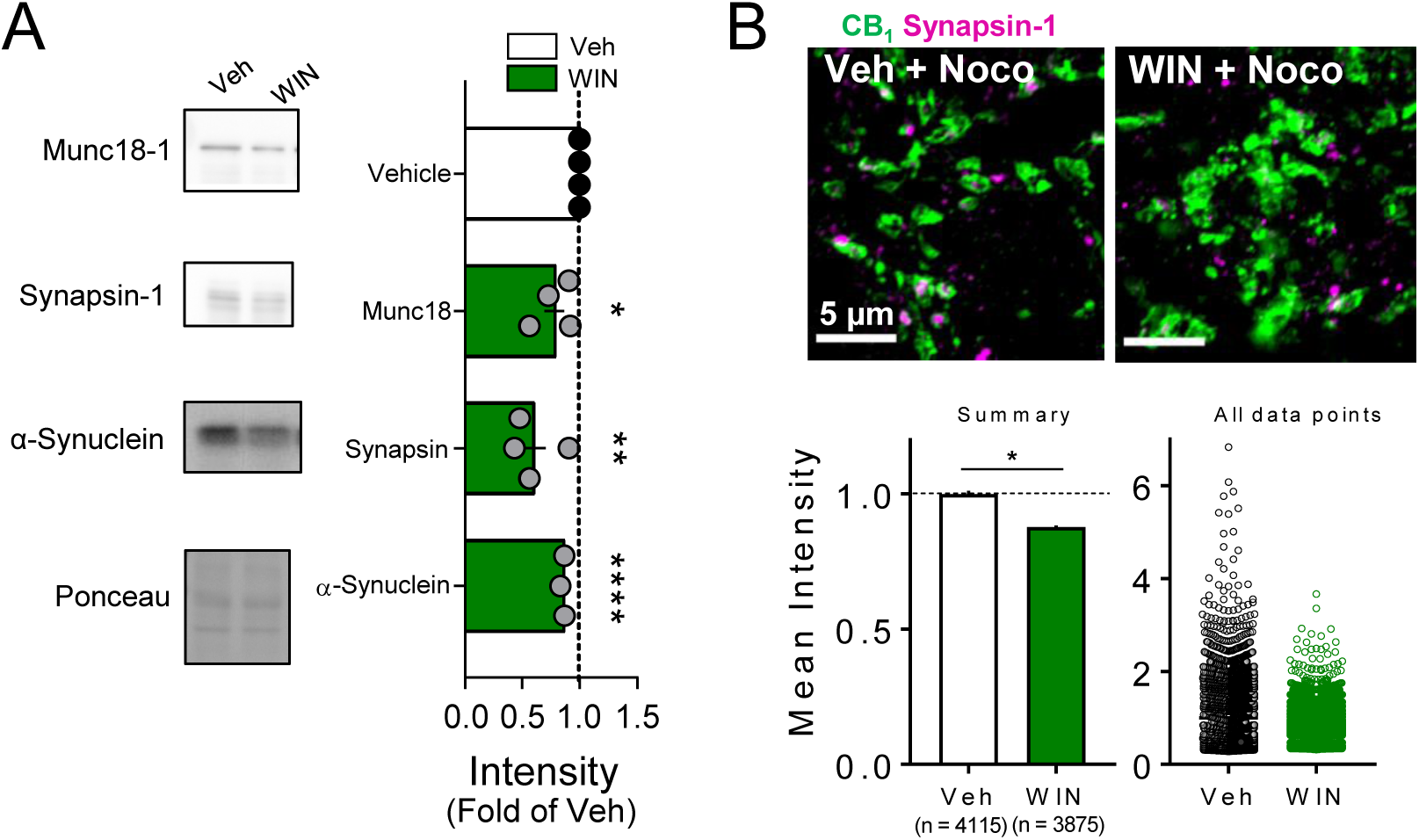
Presynaptic proteins are rapidly reduced following CB_1_ activation. *A. Left*, representative Western blot images of staining for presynaptic proteins Munc18-1, Synapsin-1, and α-Synuclein and Ponceau loading control in vehicle vs. WIN treated hippocampal cultures (5 µM, 25 min). *Right*, quantification of 3 experimental replicates normalized to Vehicle revealed a decrease in all 3 proteins consistent with SILAC. Munc18-1: 0.78 ± 0.09, p < 0.05; Synapsin-1: 0.60 ± 0.11, p < 0.01; α-Synuclein: 0.86 ± 0.01, p < 0.0001, unpaired t-test, n = number of cultures. *B. Top*, Airyscan confocal representative images of CB_1_^+^ boutons in acute hippocampal slices in CA1 pyramidal layer showing colocalization of CB_1_^+^ boutons (green) and Synapsin-1 (magenta). *Bottom left*, Summary plot (Mean ± S.E.M.) of the mean intensity of Synapsin-1 puncta within CB_1_^+^ boutons normalized to control was significantly diminished by WIN application (5 µM, 25 min). Control: 1.0 ± 0.01, WIN: 0.87 ± 0.006, Z = −2.70, p < 0.05, Mann-Whitney test, n = number of puncta analyzed (∼2 images/slice, 3 slices/rat, 4 rats/condition). *Bottom right*, plot showing distribution of all data points included in analysis.

Next, we assessed the overall contribution of the ubiquitin/proteasome system (UPS) to CB_1_-iLTD. First, to dynamically assess the UPS pathway, we measured K48-linked ubiquitinated proteins, the canonical form of ubiquitin linkage (Dantuma & Bott, 2014), following induction of CB_1_-iLTD in acute rat hippocampal slices in presence or absence of the specific proteasome inhibitor, MG-132. We found that both net flux, i.e. the amount of ubiquitinated proteins degraded by the proteasome, and the rate of degradation, measured by the ratio between blocked and basal conditions, were significantly increased. These results suggest both a larger pool of protein to degrade as well as a faster turnover rate (**Figure 5A**). However, to our surprise, CB_1-_iLTD was unaffected by application of the proteasomal inhibitor MG-132 (5 µM) during the baseline and induction (**Figure 5B**). MG-132 alone had no lasting effect on basal transmission either (**Supp. Figure 5A**). As a positive control, MG-132 application in interleaved slices resulted in accumulation of ubiquitinated proteins (**Supp. Figure 5B**). Therefore, while UPS activity is increased downstream of CB_1_ activation, proteasomal degradation is not necessary for CB_1_-iLTD.

**Figure 5.**
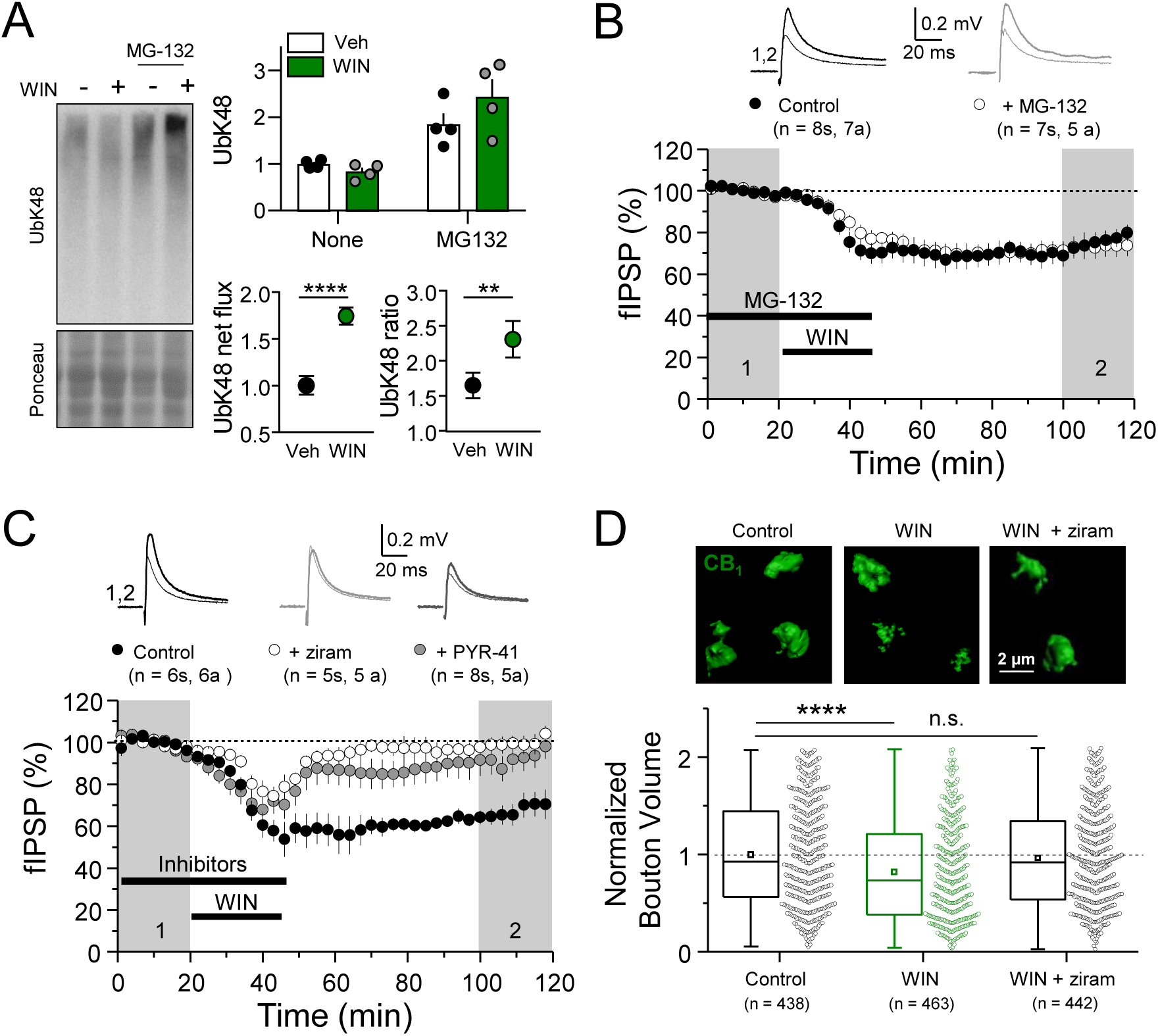
CB_1_-iLTD requires ubiquitination, but not degradation by the proteasome. *A. Left*, representative Western blot images of K48 polyubiquitin in hippocampal slices treated with Veh v. WIN or Veh v. WIN and MG-132. *Right, top*: Levels of K48 polyubiquitin following CB_1_ activation with WIN (5 µM, 25 min). *Bottom left:* UPS net flux [difference between basal (none) and proteasome blockade (MG-132) condition] is significantly increased upon CB_1_ activation. Control: 1.0 ± 0.04 vs. WIN: 1.74 ± 0.04, unpaired t-test, **** = p < 0.0001, n = 5 animals. *Bottom right*: UPS rate [ratio between MG-132 and basal condition] of K48 polyubiquitinated protein degradation is significantly increased after WIN. Control: 1.65 ± 0.08 vs. WIN: 2.31 ± 0.12, unpaired t-test, ** = p < 0.01, n = 5 animals. B. Blockade of the proteasome by bath application of MG-132 (5 µM) had no effect on iLTD. Control: 77.6 ± 4 % vs. MG-132: 73.3 ± 5 %; p > 0.05, unpaired t-test. For all electrophysiology figures, averaged summary data expressed as normalized change from baseline ± S.E.M. and n = number of slices (s), number of animals (a). C. Inhibiting ubiquitination with ziram (25 µM) or PYR-41 (50 µM) fully blocked iLTD. Control: 66 ± 5 vs. ziram: 99 ± 4 vs. PYR-41: 93 ± 6 (n = 8s,; F[2,19] = 10.22; p < 0.05, One-Way ANOVA. D. Blockade of E1 ubiquitin ligase function with ziram (25 µM, 25 min) rescued the volumetric decrease associated with CB_1_ activation by WIN (5 µM, 25 min). *Top*, representative inhibitory boutons immunolabeled with CB_1_ and reconstructed in 3D. *Bottom*, quantification of normalized CB_1_ bouton volume. Vehicle: 1.00 ± 0.03 vs. WIN: 0.823 ± 0.02 vs. WIN + ziram: 0.96 ± 0.02; F[2,1342] = 15.31; **** = p < 0.0001, One-Way ANOVA. n = number of boutons (3 images/slice, 3 slices/rat, 3 rats/condition).

Ubiquitination marks not only targets proteins for degradation, but can also affect their localization and function (Hamilton & Zito, 2013). We analyzed ubiquitination sites on a subset of proteins that were decreased by CB_1_ activation and found that most ubiquitination sites (∼60%) were located in protein-protein or protein-membrane interaction domains (**Supp. Figure 5C**) (Akimov et al., 2018), indicating that ubiquitination of these proteins could impact their function. We hypothesized that perhaps protein ubiquitination itself, independent of degradation, may play a role in CB_1_-iLTD. Using two structurally and mechanistically distinct E1 Ubiquitin ligase inhibitors, ziram and PYR-41 (Rinetti & Schweizer, 2010), we directly tested whether ubiquitination was required for CB_1-_iLTD, and found that bath application of ziram or PYR-41 blocked CB_1_-iLTD (**Figure 5C**) but had no significant effect on basal transmission (**Supp. Figure 5D**). Moreover, inhibition of ubiquitination also blocked the CB_1_-mediated decrease in CB_1_^+^ bouton volume (**Figure 5D**). These data strongly suggest that, while protein degradation follows CB_1_ activation, only protein ubiquitination is required for structural and functional CB_1-_iLTD.

## DISCUSSION

We discovered that CB_1_-iLTD involves structural changes of the presynaptic bouton that require protein synthesis. We identified the proteins that are synthesized and degraded following CB_1_ activation. Increased proteins are implicated in protein synthesis, processing and degradation, whereas decreased proteins are implicated in presynaptic structure, including Arp2/3, and function. CB_1_-iLTD involved actin remodeling, Rac1 and Arp2/3 signaling. Unexpectedly, we found that protein ubiquitination, but not proteasomal degradation, is responsible for structural and functional CB_1_-iLTD. Together, these findings point to a mechanism involving coordinated engagement of multiple cellular processes by which CB_1_^+^ presynapses can control their strength in response to CB_1_ activation.

### Presynaptic structural changes in CB_1_-iLTD

While structural changes are part and parcel of postsynaptic forms of plasticity (Bramham, 2008; Nakahata & Yasuda, 2018), and changes associated with plasticity are thought to be coordinated across the synaptic cleft, the involvement of structural changes of the presynaptic terminal in forms of long-term presynaptic plasticity are less clear. Here, we provide evidence for long-term structural changes at mature CB_1_^+^ terminals associated with CB_1_-iLTD. Previous work showed CB_1_ receptor activation can trigger ultrastructural changes in vesicle distribution associated with short-term CB_1_-mediated plasticity (Garcia-Morales et al., 2015; Ramirez-Franco et al., 2014), collapse of axonal growth cones (Berghuis et al., 2007), and inhibitory bouton formation in response to strong postsynaptic excitation (Hu et al., 2019). Our data show that transient activation of CB_1_ receptors leads to a long-term reduction of the presynaptic CB_1_^+^ compartment volume in mature somatic synapses onto CA1 pyramidal cells. Our findings (**Figure 3**) are consistent with a previous study showing that CB_1_ receptors regulate actin dynamics in growth cones by directly interacting with Rac1, a Rho GTPase (Mattheus et al., 2016), and the WAVE1 complex (Njoo et al., 2015). Other studies have suggested alternative signaling pathways by which structural changes occur downstream of CB_1_ activation. For instance, atypical coupling of CB_1_ to G_12/13_ proteins reportedly engages Rho-associated kinase (ROCK) signaling to the actomyosin cytoskeleton (Roland et al., 2014), whereas in another example, β-integrin signaling to ROCK has been implicated in a cannabinoid-mediated form of LTP (W. Wang, Jia, et al., 2018). Therefore, the signaling pathways and structural changes involved downstream of CB_1_ receptors seem to be synapse and cell type-dependent (Steindel et al., 2013). Whether CB-LTD at other synapses shares similar mechanisms remains unclear.

We showed that CB_1_-iLTD involves a rapid reduction in the size of CB_1_ boutons and the volume of active zones that requires both actin depolymerization and protein synthesis (**Figure 1**). It is unlikely that CB_1_ internalization can account for the volumetric change because our manipulation involves relatively brief WIN treatment (25 min) and WIN-induced shrinkage is evident both 25 min and 1 hour after WIN exposure, a point at which internalized receptors should be recycled back to the cell surface (Tappe-Theodor et al., 2007). Besides, the CB_1_-mediated presynaptic shrinkage is also detectable with Bassoon labeling. mRNAs for Arp2, WAVE1, and β-actin have been detected in axonal preparations and their synthesis may be important for axon formation (Donnelly et al., 2013; Spillane et al., 2012; Wong et al., 2017). However, in CB_1_-iLTD, protein synthesis seems to mediate the change in presynaptic structure via the UPS, rather than direct synthesis of actin regulators. Presynaptic terminals may vary in their structural dynamism as a function of their readiness to alter neurotransmitter release (Gogolla et al., 2007; Qiao et al., 2016). While the functional implications of presynaptic structural plasticity are yet unclear, associated changes in neurotransmitter release may result from altered Ca^2+^ channel coupling distance with the active zone (Nakamura et al., 2015), changes in numbers of docked/primed vesicles (Garcia-Morales et al., 2015; Ramirez-Franco et al., 2014), or reorganization of transsynaptic nanocolumns (Chen et al., 2018).

### Protein synthesis in presynaptic CB1-iLTD likely regulates diverse cellular processes

We have recently reported that protein synthesis is required for CB_1_-iLTD (Younts et al., 2016). Using a well-established unbiased proteomics approach in primary hippocampal neuron cultures (**Figure 2, Supp. Figure 2**) (Jordan et al., 2006; G. Zhang et al., 2011; G. Zhang et al., 2012), we identified what proteins are synthesized upon CB_1_ activation and thus can mediate CB_1_-iLTD. Despite the fact that we used whole neuron lysates, our data revealed results highly consistent with other studies that isolated presynaptic mRNAs (Bigler et al., 2017; Hafner et al., 2019; Ostroff et al., 2019; Shigeoka et al., 2016), specifically, an enrichment of mRNAs encoding for initiation factors and ribosomal proteins. We found that CB_1_ activation significantly upregulated proteins involved in protein synthesis and processing pathways (**Figure 2**). This increase in initiation factors and ribosomal proteins suggests that plasticity likely triggers an enhanced translational capacity which is consistent with our previous findings using Fluorescent Noncanonical Amino acid Tagging (FUNCAT) (Younts et al., 2016). However, why ribosomal proteins would be synthesized locally if ribosomes are exclusively assembled in the nucleolus is unclear. It is possible that in remote compartments such as axons and dendrites, damaged ribosomes could be repaired locally instead of shuttled long distances. Translation of ribosomal components could also be a novel mechanism by which local translation is regulated (Bigler et al., 2017). Surprisingly, we also noted synthesis of proteins involved in protein ubiquitination and protein degradation, which may at least partially explain the mechanism of presynaptic plasticity.

Our results have not allowed us to identify a particular protein that is being synthesized to mediate CB_1_-iLTD, but rather are suggestive of a holistic mechanism wherein CB_1_ activation impacts multiple cellular processes in concert to alter neurotransmitter release in the long-term. Although such a mechanism requires greater coordination, it would also reduce energy expenditure over time, i.e. if a presynaptic terminal will not be releasing neurotransmitter for an extended period of time (hours to days) it makes sense to degrade and recycle the release machinery, to reduce energy production, and to shrink the terminal to make space for new growth. Protein synthesis is likely necessary for the coordination and engagement of these structural, metabolic, and degradative processes. We measured changes in the protein landscape that occur fairly rapidly after CB_1_-iLTD induction (25 min) given that CB_1_-iLTD was dependent on protein synthesis during this time window (Younts et al., 2016). It is likely that additional ‘plasticity-related’ proteins are synthesized or degraded in the hours that follow iLTD induction.

### Protein degradation and presynaptic function

Proteomic analysis revealed a population of downregulated proteins involved in presynaptic function and structure, as well as energy metabolism (**Figures 1 and 4A**). In contrast, components of the protein degradation pathway, including proteasomal subunits, E2 ubiquitin ligases, and degradative enzymes, were upregulated perhaps reflecting on-demand synthesis which could regulate fast, local presynaptic protein degradation. The tight coupling of translation and degradation in the context of synaptic plasticity was described previously (Klein et al., 2015), and is presumed to occur widely in the brain as a means of maintaining proteostasis over the course of plastic changes (Biever et al., 2019; Dong et al., 2008; Hanus & Schuman, 2013; Y. C. Wang et al., 2017). This rapid activity-dependent degradation could be mediated by the recently discovered neuron-specific proteasome complex (NMP) (Ramachandran et al., 2018), although this complex is believed to target non-ubiquitinated substrates. Protein degradation has been shown to regulate presynaptic silencing, specifically by degradation of presynaptic proteins such as Rim1 and Munc-13 (Jiang et al., 2010) and CB_1_-mediated suppression of transmission at excitatory synapses via degradation of Munc18 (Schmitz et al., 2016; W. Wang, Jia, et al., 2018). However, our data strongly support the idea that protein degradation by the UPS is not directly required for CB_1-_iLTD, but does occur quickly after CB_1_ activation, as indicated by rapid loss of presynaptic proteins measured with both SILAC and western blot, presumably as a consequence of enhanced UPS activity (**Figure 4**).

We demonstrated that ubiquitination is required for CB_1_-iLTD, likely by controlling the trafficking, interactions, or the activity of its substrates, upstream of degradation (Hamilton & Zito, 2013). A previous study showed that inhibition of protein ubiquitination and degradation increased miniature EPSCs/IPSCs in cultured neurons, suggesting an important role for these processes in maintaining normal neurotransmitter release (Rinetti & Schweizer, 2010). However, in our hands, proteasomal inhibitor MG-132 had no significant effects on basal synaptic transmission and neither did E1 ubiquitin ligase inhibitors, ziram and PYR-41. To our knowledge, our study is the first to describe a mechanism of long-term presynaptic structural and functional plasticity that relies on ubiquitination. The regulation of presynaptic ubiquitination is likely achieved through the targeted expression of different E2 and E3 ubiquitin ligases (Hallengren et al., 2013) or via presynaptic cytomatrix proteins themselves (Ivanova et al., 2016; Waites et al., 2013). This raises the yet unexplored possibility that presynaptic structural dynamics and UPS activity are tightly linked.

### Potential relevance in the normal and diseased brain

CB_1_ activation via eCBs, as well as exogenous cannabinoids like Δ^9^-tetrahydrocannabinol (THC), the primary psychoactive ingredient in marijuana, can influence cognition, goal-directed behaviors, sensory processing and other critical brain functions (Araque et al., 2017; Augustin & Lovinger, 2018; Haring et al., 2012; Heifets & Castillo, 2009; Hoffman & Lupica, 2013; Zlebnik & Cheer, 2016). Cannabinoid signaling has also been implicated in several brain disorders (Zou & Kumar, 2018). Autism is broadly associated with changes in synaptic protein levels, but also disruption of CB_1_-LTD (Busquets-Garcia et al., 2014; Chakrabarti et al., 2015). In particular, in a mouse model of Fragile X Syndrome (FXS), the most common monogenic cause of autism, where RNA-binding protein, Fragile X mental retardation protein (FMRP) is deleted (Bagni & Zukin, 2019), eCB-mediated plasticity in the hippocampus, striatum, prefrontal cortex is impaired (Jung et al., 2012; Maccarrone et al., 2010; Martin et al., 2017; W. Wang, Cox, et al., 2018; L. Zhang & Alger, 2010). Although changes in the eCB mobilization in FXS may explain some of the impairment (Jung et al., 2012; Maccarrone et al., 2010; L. Zhang & Alger, 2010), the role of FMRP in the regulation of local presynaptic protein synthesis may also play a role (but see Jung et al., 2012), though this remains to be tested (Busquets-Garcia et al., 2013). Many neurodegenerative diseases are characterized by imbalanced proteostasis, including Alzheimer’s disease (AD) and Parkinson’s disease (PD), resulting in pathological accumulations of misfolded proteins (Klaips et al., 2018). WIN treatment has been shown in animal models of AD and PD to be neuroprotective and to alleviate cognitive and motor symptoms (Basavarajappa et al., 2017), potentially through the ability of the CB_1_ receptor to regulate synaptic proteostasis.

## METHODS

### Immunohistochemistry and Microscopy

Acute rat hippocampal slices were made as described below for electrophysiological recordings and allowed to recover for at least 1 hour after slicing. Slices were incubated in beakers containing ACSF and drug treatments described in Results and underwent constant oxygenation. Slices were fixed immediately after treatments in 4% PFA in PBS overnight at RT. Slices were washed twice in PBS then incubated in blocking buffer (4% BSA in PBS + 0.1% Tx-100) for 1 hour at RT. Primary antibodies (CB_1_, 1:1000, Immunogenes (Budapest, Hungary); Synapsin-1 1:1000 Synaptic Systems (Goettingen, Germany) #106001; Bassoon, 1:1000, Enzo Life Sciences (Farmingdale, NY, USA); Paravalbumin, 1:1000, Sigma Aldrich) were diluted directly into the blocking buffer and floating slices were incubated overnight at 4C. After 4 washes with PBS, slices were incubated in secondary antibodies (Invitrogen) diluted in blocking buffer overnight at 4C. Slices were washed 5X with PBS, then mounted. Images were acquired on a Zeiss LSM 880 with Airyscan using a Plan-Apochromat 63×/1.4 Oil DIC M27 and 1.8X zoom. Images were Airyscan processed prior to analysis. Pixel width and height was 0.049 µm and voxel depth was 0.187 µm. Imaris 9.2 software was used to reconstruct boutons in 3D using the Surface function. Threshold, laser power, and gain were kept constant for each experiment.. CB_1_ boutons were screened after 3D reconstruction to ensure correct identification. Only boutons that fell between 0.01-2 µm^3^, did not touch the image border, and had a sphericity value above 0.3 were considered. For Bassoon (Figure 1C), a custom-written macro on FIJI was used to remove all Bassoon signal that didn’t colocalize with CB_1_ labeling, then Imaris was used to quantify the bassoon signal. FIJI was used to analyze synapsin puncta intensity (Figure 4B) inside CB_1_ boutons using a custom-written macro. All imaging and analysis was performed blind to treatment group.

### SILAC

Primary hippocampal neurons were prepared from E18-19 rat brains and grown on poly-D-lysine coated 15 cm plates at a density of 3.5 million cells/ 15 cm plate in DMEM media without l-arginine or l-lysine (Cambridge Isotope Laboratories, Tewksbury, MA, USA, Cat# DMEM-500), with pen/strep and B-27 supplement (Invitrogen, Carlsbad, CA, USA). 84mg/L of l-arginine 13C_6_ (Sigma Aldrich, St. Louis, MO, USA) and 146mg/ L of l-lysine D_4_ (Thermofisher, Waltham, MA, USA) was supplemented for ‘medium’ labelled media and 84mg/L of l-arginine 13C_6_15N_4_ and 146mg/ L of l-lysine 13C_6_15N_4_ was added to ‘heavy’ labelled media. Neurons were grown in this media for 15 days, which results in >90% incorporation of labeled amino acids into the cellular proteomes (G. Zhang et al., 2011; G. Zhang et al., 2012). For treatment, neuron cultures DIV 16 received 15 ml fresh media containing either vehicle or WIN (5 µM) for 25 min. Neurons were washed 3X with ice-cold PBS without Mg^2+^ or Ca^2+^ (0.01 M, pH = 7.4). After 3 washes, cells were harvested and lysed in SDS lysis buffer containing: (50 mM Tris, 2% SDS, 2mM EDTA) for 30 min at RT. Lysates were then sonicated briefly, allowed to incubate for another 30 min at RT, and centrifuged for 5 min at 15,000g to remove insoluble debris. 10 µg of lysate from ‘medium’ cells treated with vehicle were mixed with 10 µg ‘heavy’ WIN treated cells (Forward sample). Separately, 10 µg of lysate from ‘medium’ cells treated with WIN were mixed with 10 µg ‘heavy’ vehicle treated cells (Reverse sample). These mixtures (20 µg total protein) were loaded onto a 10% Bis-Tris gel and subjected to SDS-PAGE. The gel was stained with Coomasie for 1 hr and protein lanes were cut up into 12 equal sized portion in order to improve protein coverage (Jordan 2004). Mass spectrometry was performed in collaboration with the Einstein Proteomic Facility using the LTQ-Orbitrap hybrid mass spectrometer (Thermo Fisher Scientific) equipped with a nanoelectrospray ionization source (Jamie Hill Instrument Services) was used. An Eksigent nanoLC system (Eksigent Technologies, Dublin, CA, USA) equipped with a self-packed 750-μm × 12-cm reverse phase column (Reprosil C18, 3 μm, Dr. Maisch, Germany) was coupled to the mass spectrometer. Peptides were eluted by a gradient of 3–40% acetonitrile in 0.1% formic acid over 110 min at a flow rate of 300 nL/min. Mass spectra were acquired in data-dependent mode with one 60,000 resolution MS survey scan by the Orbitrap and up to eight MS/MS scans in the LTQ for the most intense peaks selected from each survey scan. Automatic gain control target value was set to 1,000,000 for Orbitrap survey scans and 5,000 for LTQ MS/MS scans. Survey scans were acquired in profile mode and MS/MS scans were acquired in centroid mode. Charge state deconvolution and deisotoping were not performed. All MS/MS samples were analyzed using Mascot (Matrix Science, London, UK; version 2.5.1.3). Mascot was set up to search the NCBInr_20160618.fasta; SwissProt_AC_20170620 database (selected for Rattus, 92376 entries) assuming the digestion enzyme stricttrypsin. Mascot was searched with a fragment ion mass tolerance of 0.40 Da and a parent ion tolerance of 50 PPM. Carbamidomethyl of cysteine was specified in Mascot as a fixed modification. Deamidated of asparagine and glutamine, label:2H(4) of lysine, label:13C(6) of arginine, label:13C(6)15N(2) of lysine, label:13C(6)15N(4) of arginine and oxidation of methionine were specified in Mascot as variable modifications. Scaffold (version Scaffold_4.8.3, Proteome Software Inc., Portland, OR) was used to validate MS/MS based peptide and protein identifications. Peptide identifications were accepted if they could be established at greater than 95.0% probability by the Scaffold Local FDR algorithm. Protein identifications were accepted if they could be established at greater than 99.0% probability and contained at least 2 identified peptides. Protein probabilities were assigned by the Protein Prophet algorithm (Nesvizhskii, Al et al Anal. Chem. 2003;75(17):4646-58). Proteins that contained similar peptides and could not be differentiated based on MS/MS analysis alone were grouped to satisfy the principles of parsimony. Proteins sharing significant peptide evidence were grouped into clusters.

### Gene ontology analysis

SILAC results were ranked according based on fold change and submitted to a GSEA Preranked analysis in GSEA (v. 4.0.2) with 1000 permutations. Terms smaller than 15 genes or bigger than 500 were discarded as previously reported (Merico et al., 2010). The enrichment map was generated in Cytoscape (3.7.1) (Kucera et al., 2016) using Enrichment map plugin (3.2.0) (Merico et al., 2010) using the following thresholds: p value <0.05, FDR <0.001. The overlap coefficient was set to 0.5. For confirmation, we also performed Gene Ontology analysis using two other tools. First, filtered lists (|log_2_ fold change| > 0.5) were analyzed through the use of IPA (QIAGEN Inc., Hilden, Germany, https://www.qiagenbioinformatics.com/products/ingenuitypathway-analysis). Then, we performed ontology enrichment using a recently published expert-curated knowledge database for synapses (Koopmans et al., 2019). Terms were selected with a FDR<0.01. The parental term ‘Synapse’ was discarded as not being informative (e.g. to general).

### Electrophysiology Slice Preparation and Recording

Experimental procedures adhered to NIH and Albert Einstein College of Medicine Institutional Animal Care and Use Committee guidelines. Acute transverse slices were prepared from young adult male and female Sprague Dawley rats (P18-27). The cutting solution contained (in mM): 215 sucrose, 20 glucose, 26 NaHCO_3_, 4 MgCl_2_, 4 MgSO_4_, 1.6 NaH_2_PO_4_, 2.5 KCl, and 1 CaCl_2_. The artificial cerebral spinal fluid (ACSF) recording solution contained (in mM): 124 NaCl, 26 NaHCO_3_, 10 glucose, 2.5 KCl, 1 NaH_2_PO_4_, 2.5 CaCl_2_, and 1.3 MgSO_4_. After ice-cold cutting, slices recovered at RT (in 50% cutting solution, 50% ACSF) for <30 min and then at room temperature (RT) for 1 hr in ACSF. All solutions were bubbled with 95% O_2_ and 5% CO_2_ for at least 30 min. Although the form of long-term inhibitory synaptic plasticity studied here (i.e. iLTD) is present under physiological recording conditions at 37 °C (Younts et al., 2013), inhibitory synaptic transmission is less stable at this temperature, and therefore we conducted our experiments at 25.5 ± 0.1°C.

For extracellular field recordings, a single borosilicate glass stimulating pipette filled with ACSF and a glass recording pipette filled with 1M NaCl were placed approximately 100 µm apart in *stratum pyramidale*. To elicit synaptic responses, paired, monopolar square-wave voltage or current pulses (100–200 μs pulse width) were delivered through a stimulus isolator (Isoflex, AMPI) connected to a broken tip (∼10–20 μm) stimulating patch-type micropipette filled with ACSF. Typically, stimulating pipettes were placed in CA1 stratum pyramidale (150–300 μm from the putative apical dendrite of the recorded pyramidal cell, 150–200 μm slice depth). Stimulus intensity was adjusted to give comparable magnitude synaptic responses across experiments less than ∼0.6 mV). Inhibitory synaptic transmission was monitored in the continuous presence of the NMDA receptor antagonist d-(-)-2-amino-5-phosphonopentanoic acid (d-APV; 25 μM), the AMPA/kainate receptor antagonist 2,3-dihydroxy-6-nitro-7-sulfonyl-benzo[f]quinoxaline (NBQX; 5 μM), and the μ-opioid receptor agonist, [D-Ala^2^, N-MePhe^4^, Gly-ol]-enkephalin (DAMGO, 50 nM). To elicit chemical-iLTD (Heifets et al., 2008) the CB_1_ agonist WIN 55,212-2 (WIN; 5 µM) was bath applied for 25 min, and 5 stimuli at 10 Hz were delivered at 0.1 Hz during the last 10 min of WIN. WIN was chased with the CB_1_ inverse agonist/antagonist SR 141716 (5 µM) or AM251 (5 µM) to halt CB_1_ signaling. Baseline and post-induction synaptic responses were monitored at 0.05 Hz during iLTD. Stimulation and acquisition were controlled with IgorPro 7 (Wavemetrics). Shaded boxes in figures correspond to when plasticity was analyzed with respect to baseline and when representative traces were collected and averaged. Summary data (i.e. time-course plots and bar graphs) are presented as mean ± standard error of mean (SEM). PPR was defined as the ratio of the amplitude of the second EPSC (baseline taken 1-2 ms before the stimulus artifact) to the amplitude of the first EPSC. The magnitude of LTD was determined by comparing 20 min baseline responses with responses 80-100 min post-LTD induction.

### Western Blotting

Protein concentration was determined using the Lowry method with bovine serum albumin as a standard (Lowry et al., 1951). Primary hippocampal neurons or hippocampal slices were solubilized on ice with RIPA buffer (1% Triton X-100, 1% sodiumdeoxycholate, 0.1% SDS, 0.15MNaCl, 0.01Msodium phosphate, pH7.2) followed by sonication. Immunoblotting was performed after transferring SDS-PAGE gels to nitrocellulose membrane and blocking with 5% low-fat milk for 1h at room temperature. The proteins of interest were visualized after incubation with primary antibodies (α-synuclein 1:1000 BD Biosciences (San Jose, CA, USA) #610787; Synapsin-1 1:1000 Synaptic System (Goettingen, Germany) #106001; Munc-18-1 1:1000 Synaptic System #1160002; Arp2/3 1:1000 Novus Biologicals (Centennial, CO, USA) # NBP188852; Ubiquitin K48 1:1000 EMD Millipore (Burlington, MA, USA) #05-1307; Ubiquitin K63 1:1000 EMD Millipore #05-1308) by chemiluminescence using peroxidase-conjugated secondary antibodies in LAS-3000 Imaging System (Fujifilm, Tokyo, Japan). Densitometric quantification of the immunoblotted membranes was performed using ImageJ (NIH). All protein quantifications were done upon normalization of protein levels to Ponceau staining. Ponceau normalization was chosen over comparison to actin as our work and others showed that CB_1_-iLTD induces modification of actin cytoskeleton.

### Ubiquitination sites analysis

Ubiquitination sites were identified using Ubisite, a publicly available resource for ubiquitination site prediction (Akimov et al., 2018). To minimize false positive rate, confidence level was set on high. When available, functional domains were annotated using the uniport.org database (UniProt, 2019).

### Data analysis, Statistics and Graphing

Analysis and statistics were carried out in OriginPro 2015 (OriginLab) and Graphpad Prism 7.02. Significance (p <0.05) was assessed with one-way ANOVA (means comparison with *post hoc* Bonferroni test), Student’s paired and unpaired t-tests, Wilcoxon matched-pairs signed rank test, Mann Whitney U test, or Pearson’s correlation coefficient, as indicated. All experiments were performed in an interleaved fashion. Unless stated otherwise, “n” represents number of field recordings in slices. All experiments include at 3 animals. Plotting of SynGO results was made using matplotlib (3.0.3)(Hunter, 2007) in Python (3.7. 3)(Oliphant, 2007) environment.

### Reagents

Stock reagents were prepared according to the manufacturer’s recommendation in water, DMSO (<0.01% final volume during experiments), or phosphate buffered saline (PBS), stored at −20°C, and diluted into ACSF or intracellular recording solutions as needed. CNQX, D-APV, SR 141716, and WIN were acquired from the NIMH Chemical Synthesis and Drug Supply Program; salts for making cutting, ACSF, ziram, and intracellular recording solutions from Sigma Aldrich (St. Louis, MO, USA); AM251, NSC-23766, MG-132, DAMGO, cycloheximide from Tocris Bioscience (Bristol UK); jasplakinolide, anisomycin, PYR41 from Cayman Chemical (Ann Arbor, MI, USA); CK-666 from EMD Millipore. Reagents were either acutely bath applied, diluted into the intracellular recording solution, or preincubated with slices/cultures, as indicated in Results.

## Acknowledgements

We thank Dr. Thomas Younts, Dr. Matthew Klein, and Dr. Ana Maria Cuervo for helpful discussions. We thank Dr. Kostantin Dobrenis, Kevin Fisher, and Vladimir Mudragel of the Einstein Neural Cell Engineering and Imaging Core for their advice and assistance with Airyscan confocal microscopy acquisition and analysis. We thank Edward Nieves of the Einstein Proteomics Facility for assistance performing and analyzing proteomic data. This research was supported by the National Institutes of Health: F31MH114431 to HRM, R01-MH081935, R01-DA17392 and R01-NS113600 to PEC, a shared instrument grant (1S10OD25295) to KD, R01-AG039521 to BAJ, MB was supported by the Rainwater Charitable Foundation. HM and MB also received Junior Investigator Neuroscience Research Award (JINRA) from Dominick P. Purpura Department of Neuroscience at Albert Einstein College of Medicine for this project.

The authors declare no competing financial interests.

## Author Contributions

H.R.M. and P.E.C wrote the manuscript. H.R.M., M.B., B.A.J., A.M.C., and P.E.C designed experiments. H.R.M. performed SILAC, electrophysiology, and imaging experiments. M.B. analyzed SILAC and performed western blots.

---

**Supplemental Figure 1 (related to Figure 1):**
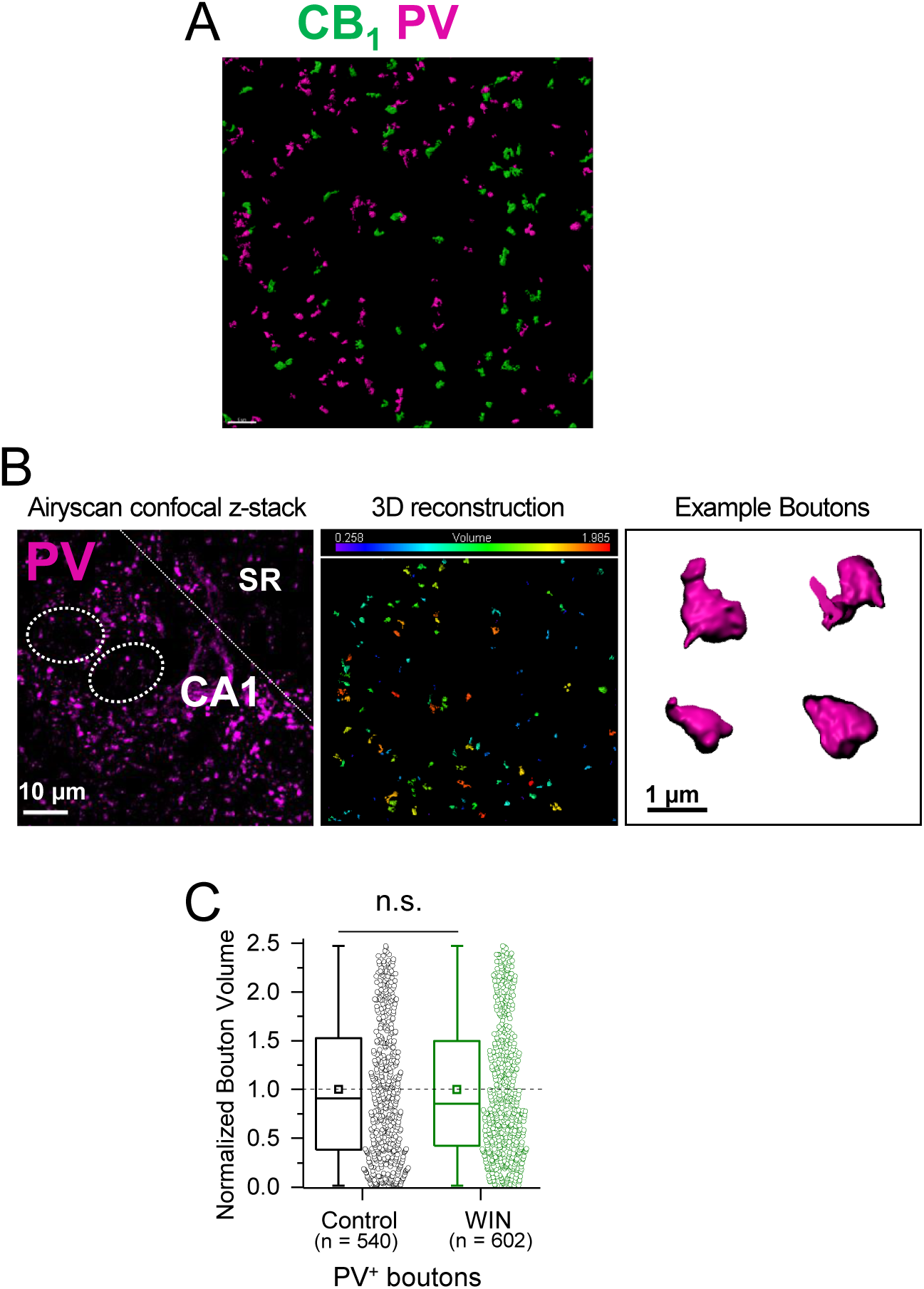
A. Representative widefield confocal maximum projection of CA1 pyramidal layer showing PV and CB_1_^+^ boutons. *B. Left*, representative widefield confocal z-stack of CA1 pyramidal layer and 3D Imaris reconstruction of CB_1_^+^ boutons. Dotted lines indicate location of putative pyramidal cell somas. *Center*, volume of individual boutons was quantified using dense PV immunolabeling of inhibitory interneuron terminals. *Right*, representative single boutons reconstructed in 3D. C. Quantification of bouton volume normalized to control. Activation of CB1 receptors with WIN (5 µM, 25 min) resulted in no change in PV bouton volume. Control: 1.0 ± 0.03 v. WIN: 0.99 ± 0.03, U = 161715.5, Z = −0.148, n.s. indicates not significantly different, Mann-Whitney, n = number of PV^+^ boutons (2 images/ slice, 2 slices/rat, 5 rats/condition). Data are presented as box plots (left) and data points (right) where box represents the 25^th^ and 75^th^ percentile of data range, mean is represented with a square, and median with a line inside the box. Minimum and maximum of data are given by the capped line.

**Supplemental Figure 2 (related to Figure 2):**
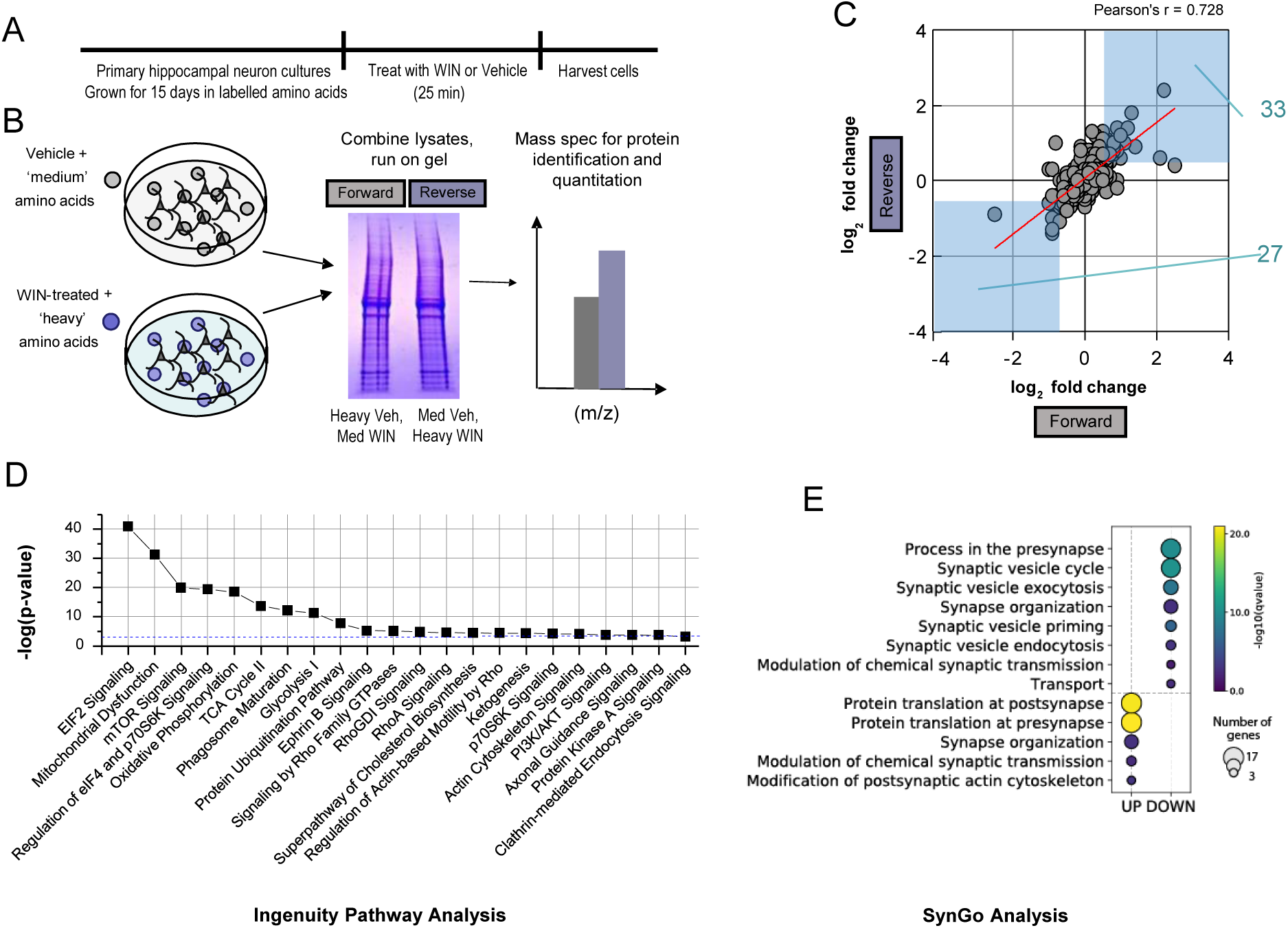
A. Experimental timeline for SILAC. Neurons were grown for 15 days in isotope-labelled amino acids, before being treated for 25 min with WIN and harvested. B. Schematic of workflow to identify proteins. Cultures were lysed and combined to give ‘forward’ and ‘reverse’ replicates. Mass spectrometry was performed for identification and quantification of proteins altered by WIN treatment. C. Scatterplot of all proteins identified in forward and reverse experiments (n = 391) by log_2_ fold change. Forward and reverse experiments are significantly correlated indicating good reproducibility (Pearson’s correlation coefficient = 0.72799, p < 0.05). Proteins that fall into light blue boxes were considered significantly regulated by WIN treatment (log_2_ fold change ≥ ±0.5). 33 total proteins were upregulated and 27 downregulated. D. Top 22 selected pathways identified by Ingenuity Pathway Analysis (IPA) ranked by −log10(p-value). Blue dashed line indicates p-value = 0.001, all pathways have p < 0.001. E. SynGO term analysis identified significant upregulation of local synaptic translation pathways and downregulation of GO terms involved in presynaptic function, as indicated by log q-value. Size of circle represents number of genes identified per SynGO term.

**Supplemental Table 1: Full protein list and Ingenuity Pathway Analysis**

**Supplemental Figure 3 (related to Figure 2):**
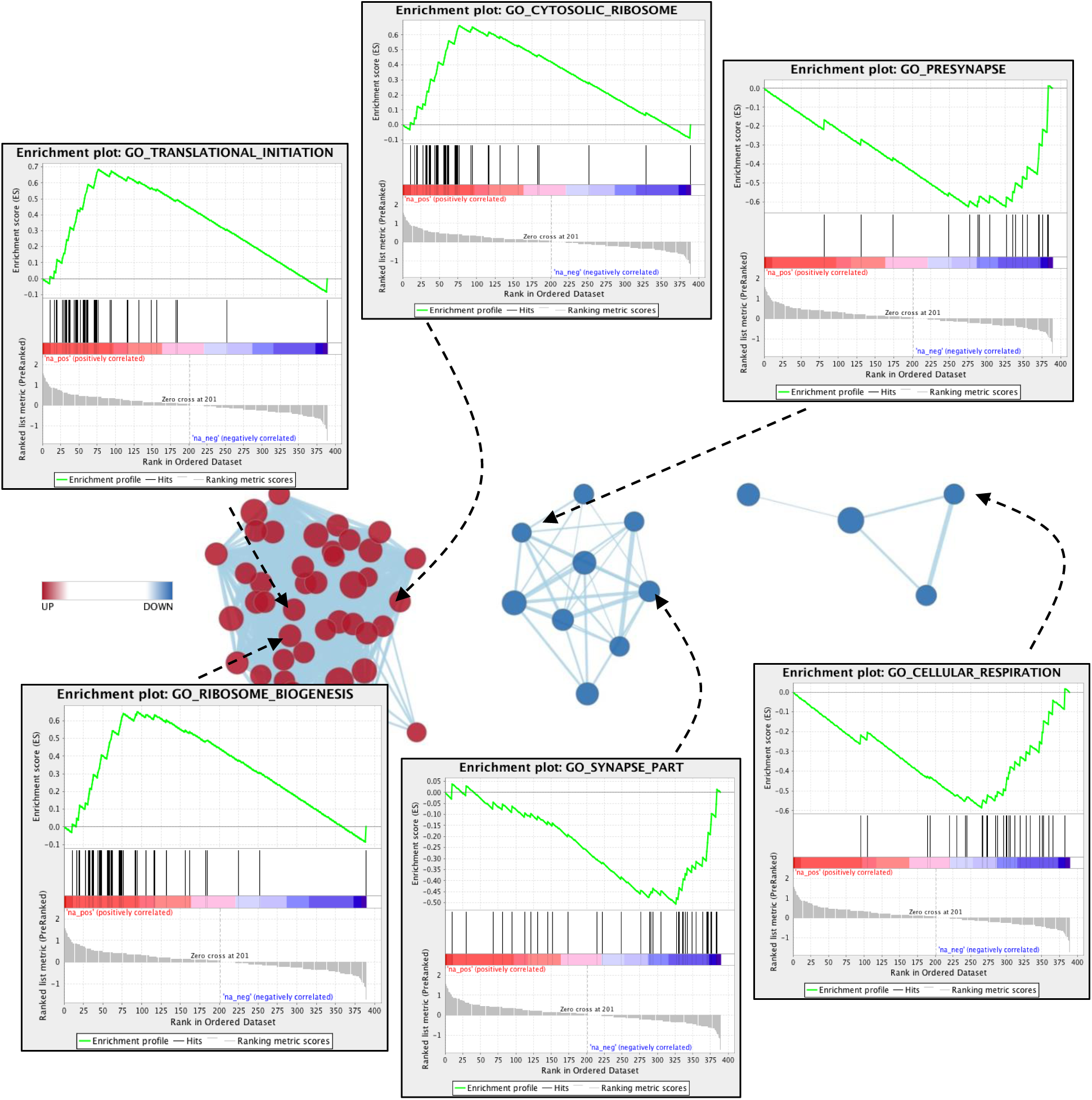
Examples of GO terms in each cluster are provided. Enrichment score (green line) is computed by individual protein rank (black vertical lines) in the ranked list for each GO term and direction of change (up v. down v. no change).

**Supplemental Table 2: Raw data for GSEA and Enrichment Map Analysis**

**Supplemental Figure 4 (related to Figure 3):**
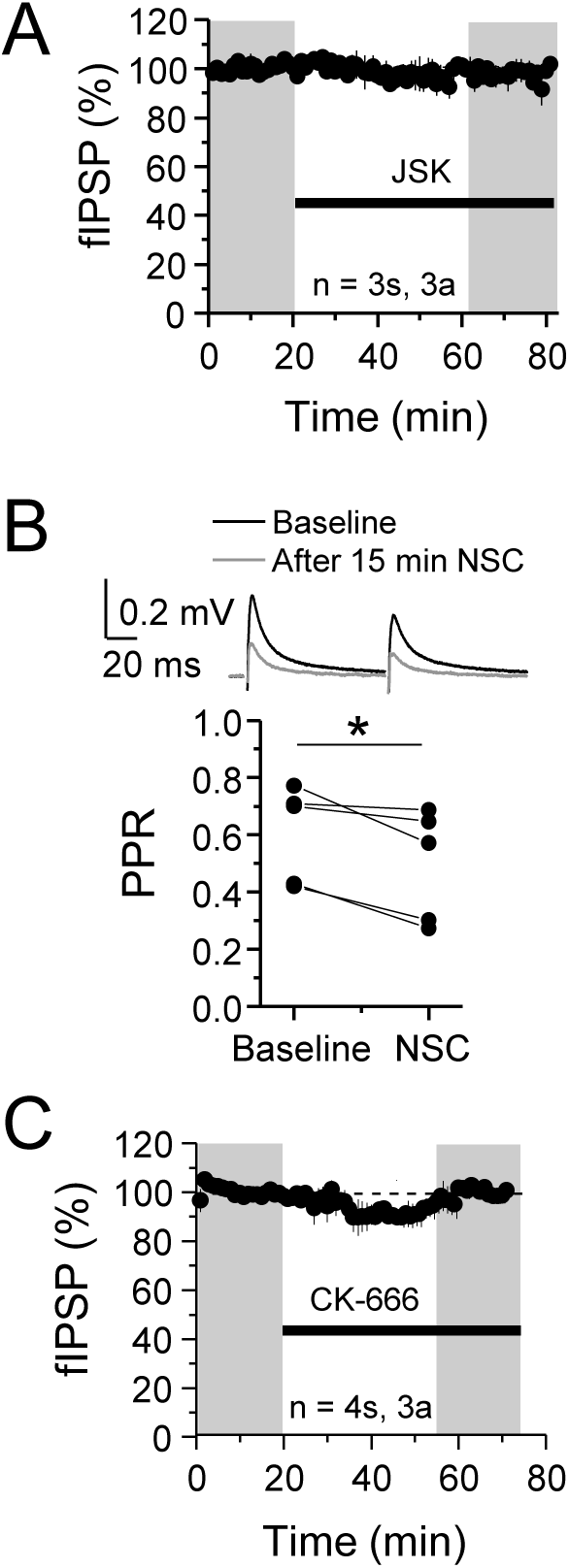
A. JSK (250 nM) had no effect on baseline. JSK: 97.76 ± 2.8, one sample t-test, p > 0.05. For all electrophysiology figures, averaged summary data expressed as normalized change from baseline ± S.E.M. and n = number of slices (s), number of animals (a). B. NSC application on baseline caused decreased PPR (measured at 35-45 min) suggesting a presynaptic target. Pre: 0.65 ± 0.06 v. NSC: 0.57 ± 0.08, paired-sample t-test, p < 0.05, n = 5 slices (same as Figure 4B). C. CK-666 (100 µM) had no effect on baseline. CK-666: 95.44 ± 5.0, one sample t-test, p > 0.05.

**Supplemental Figure 5 (related to Figure 5):**
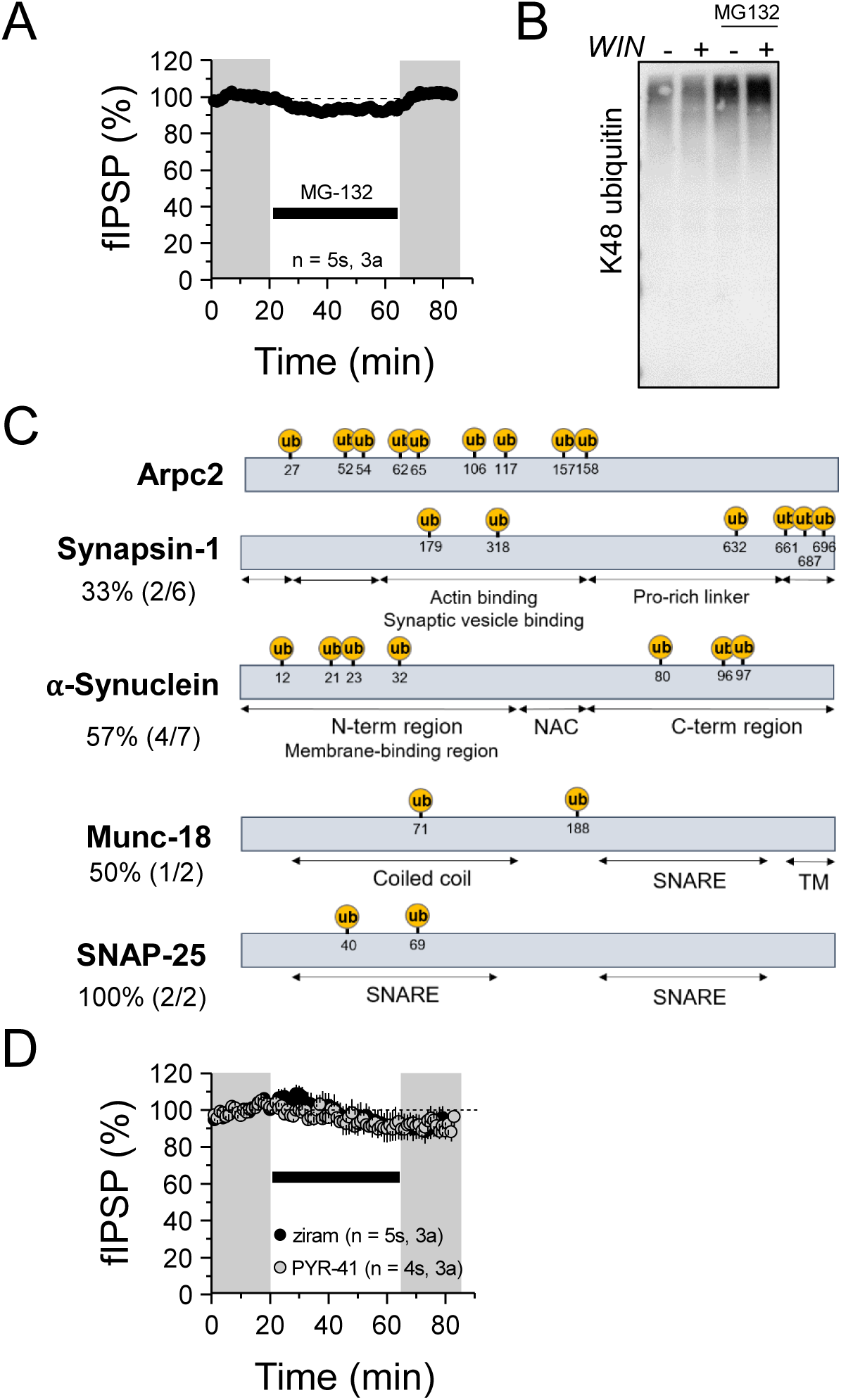
A. MG-132 (5 µM) did not cause a lasting change in baseline transmission. MG-132: 99.66 ± 2.43 %; p > 0.05, one sample t-test. For all electrophysiology figures, averaged summary data expressed as normalized change from baseline ± S.E.M. and n = number of slices (s), number of animals (a). B. Western blot of accumulated K48 ubiquitinated proteins when proteasome is blocked with 5 µM MG-132 demonstrated the efficacy of the drug. C. Predicted ubiquitination sites based on protein sequence plotted with UbiSite. Percentage represents # of predicted Ub sites in functional domains of protein divided by total predicted sites. Note: no protein structure data was available for Arpc2. NAC: non-amyloid component region of α-synuclein. SNARE: SNARE protein interaction site. TM: transmembrane domain. D. Ziram (25 µM) or PYR-41 (50 µM) did not cause a significant change in baseline transmission. ziram: 91.56 ± 6.49 %; p > 0.05, one sample t-test. PYR-41: 91.49 ± 4.0; p > 0.05, one sample t-test.

